# Anchor-Enhanced Bead Design for Reduced Oligonucleotide Synthesis Errors in Single-cell sequencing

**DOI:** 10.1101/2024.04.08.587145

**Authors:** Jianfeng Sun, Martin Philpott, Danson Loi, Gabriela Hoffman, Jonathan Robson, Neelam Mehta, Eleanor Calcutt, Vicki Gamble, Tom Brown, Tom Brown, Udo Oppermann, Adam P Cribbs

## Abstract

Single-cell transcriptomics, reliant on the incorporation of barcodes and unique molecular identifiers (UMIs) into captured polyA+ mRNA, faces a significant challenge due to synthesis errors in oligonucleotide capture sequences. These inaccuracies, which are especially problematic in long-read sequencing, impair the precise identification of sequences and result in inaccuracies in UMI deduplication. To mitigate this issue, we have modified the oligonucleotide capture design, which integrates an interposed anchor between the barcode and UMI, and a ’V’ base anchor adjacent to the polyA capture region. This configuration is devised to ensure compatibility with both short and long-read sequencing technologies, facilitating improved UMI recovery and enhanced feature detection, thereby improving the efficacy of droplet-based sequencing methods.

## Introduction

Droplet-based sequencing has rapidly advanced to be the gold standard method in studying single-cell transcriptomics by leveraging its capability to offer higher throughput. Droplet-based sequencing methods such as Drop-seq^1^, InDrops^2^ and 10X Chromium^3^ provide improved throughput at a reduced cost per cell. While Drop-seq gained traction as a favourable academic approach, the 10X Genomics Chromium platform capitalised on this momentum and commercialised droplet-based single-cell sequencing, ultimately securing its position as the market front-runner.

Central to this technology are oligonucleotide sequences that are synthesised and contain a polyA^+^ capture region, a PCR primer, a cell barcode and a unique molecular identifier (UMI). Historically, the techniques and strategies underpinning droplet-based sequencing were developed with Illumina sequencing platforms in mind. Typically, the Illumina sequencing protocol employs a fragmentation step so that sequencing of Read 1 contains both the barcode and the UMI sequence, while, Read 2 contains the sequencing information for either the 3’ or 5’ end of the captured transcript. However, the emergence of long-read sequencing, which allows for complete end-to-end sequencing, has made identification of the barcode and UMI sequences more challenging. This stems from the reliance on computational pattern matching to locate the primer site upstream of the barcode and UMI to ensure accurate identification. PCR and sequencing errors further complicate this^4, 5^, making barcode and UMI identification less accurate and leading to a significant proportion of reads being discarded.

In previous work, we introduced homodimer and homotrimer UMIs to counteract PCR and sequencing inaccuracies across both bulk and single-cell resolutions^4, 5^. In this study, we identify another critical source of error: bead oligonucleotide synthesis inaccuracies. To address this, we have developed a bead design that alleviates these synthesis issues. Crucially, our design aligns with both short-read and long-read sequencing methods. Our findings suggest that anchors incorporated into the oligonucleotide capture sequence can enhance UMI delineation. Additionally, we’ve incorporated an anchor sequence between the barcode and UMI, promoting precise and efficient UMI recognition. These adjustments have resulted in a marked improvement in UMI recovery and a heightened transcript detection rate, improving the capabilities of droplet-based sequencing.

## Results

### Evidence of bead truncation in 10X chromium and Drop-seq beads

Droplet based single-cell methods such as Drop-seq and 10X Chromium process mRNA from individual cells encapsulated in oil droplets in a highly parallel fashion. Here, cells are trapped within a droplet, which facilitate cell lysis and mRNA capture using oligonucleotide beads. However, bead designs vary between Drop-seq and 10X Chromium (**Fig. 1a**). In Drop-seq, the bead contains a PCR primer region, followed by a 12-base pair (bp) cell barcode created by a split and pool synthesis. Next is an 8bp Unique Molecular Identifier (UMI) sequence, with a V (A, C, or G) base preceding the start of a poly(dT) capture region that acts to couple the barcode and UMI to the polyA^+^ mRNA. In contrast, 10X Chromium bead design is distinct. It contains a 16bp barcode formed though combinatorial enzymatic ligation split and pool, followed by a 12bp UMI sequence. Unlike Drop-seq, the 10X Chromium beads lack a V base between the UMI and the poly(dT) sequence. They uniquely incorporate a V base followed by an N (A, C, G or T) base at the poly(dT) end, ensuring mRNA capture near the polyA terminus of the mRNA.

**Figure 1:**
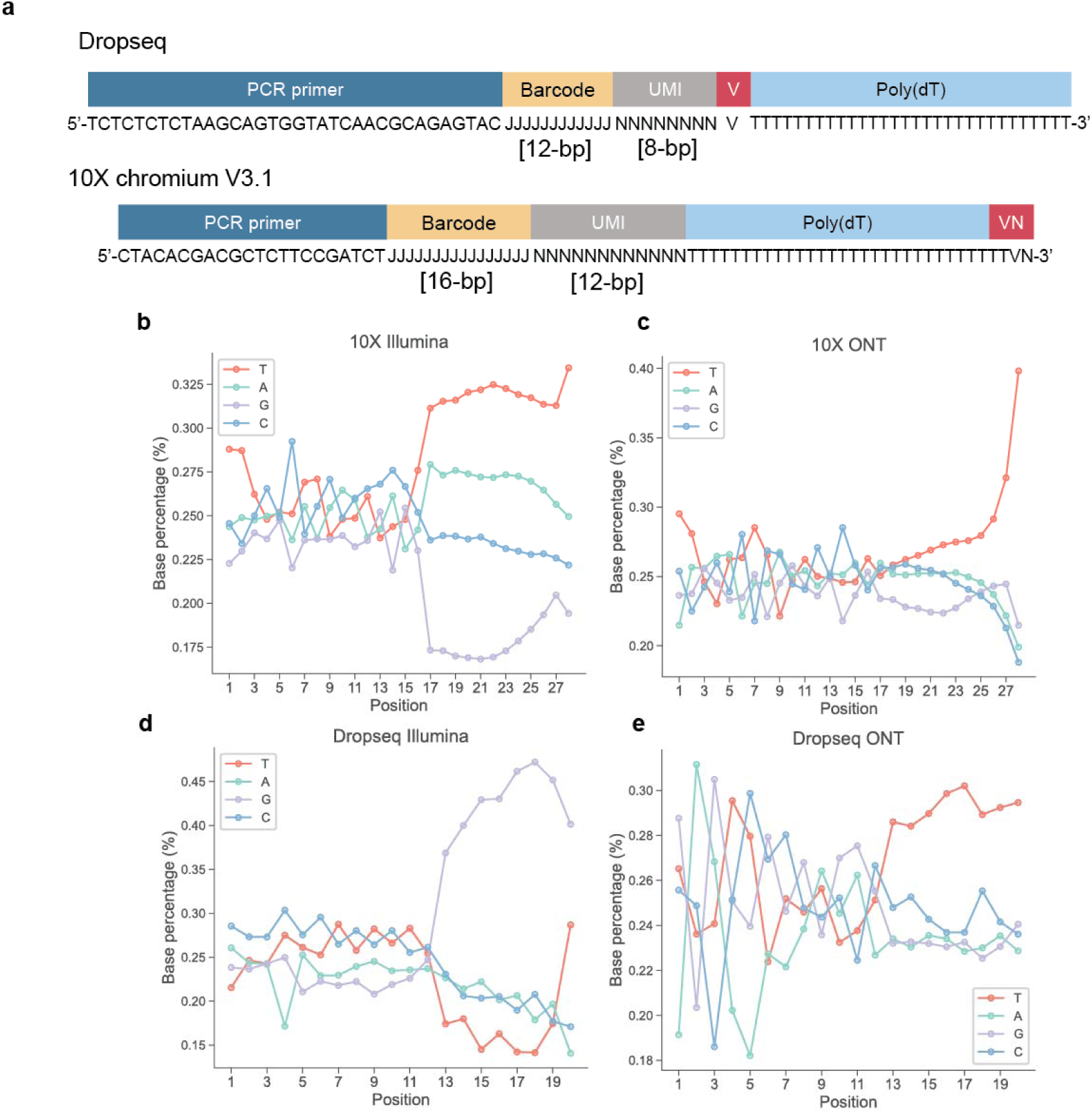
Analysis of the base composition of 10X Chromium and Drop-seq barcodes and UMIs. **a**, A comparative schematic illustrates the bead designs of both Drop-seq and 10X Chromium platform v3.1. While both designs incorporate a barcode and UMI, they vary in length, the positioning of the V base (A, C, or G), and PCR primer sequences. N signifies a random base, J denotes a semi-random base, and T, C, G and A represent Thymidine, Cytosine, Guanine, Adenosine, respectively. **b**, The 10X Chromium 5k PBMC v3.1 dataset was downloaded, and analysis was performed for short-read Illumina sequencing. **c**, The 10X Chromium 5k PBMC v3.1 dataset was downloaded, and analysis was performed for long-read ONT sequencing. **d**, Illumina sequencing data from Macosko et al 2015 was downloaded and analysed. **e**, A mixed species Drop-seq single-cell experiment was performed then sequenced using ONT’s PromethION^TM^ device. **b-e**, The proportion of each base is plotted across the barcode and UMI sequences.

High accuracy in oligonucleotide synthesis is crucial for single-cell sequencing success, essential to minimise misreads from faulty barcode matching and prevent inflated count matrices due to misidentified UMI sequences. However, despite the tremendous impact of synthetic oligonucleotides in biology, solid-phase phosphoramidite oligonucleotide synthesis is a multi-step chemical process which can never be perfect^6^. The efficiency of nucleoside phosphoramidite coupling is around 99% per cycle^7, 8^, hence the purity of full-length oligonucleotides decreases significantly, and a 100-mer oligonucleotide will typically contain less than 50% of strands with the desired sequence^9^. Synthesis errors in oligonucleotides, including substitutions, insertions, and deletions, are proportionate to their length, with capping steps reducing deletions but increasing G to A transitions^10^. Consequently, purification methods can be utilised to extract such contaminants from shorter oligonucleotide sequences. However, the unique barcode requirements for each bead precludes the application of purification methods in droplet based single-cell sequencing. As a result, synthesis errors present a significant and unresolved challenge within the field of single-cell sequencing.

To explore the occurrence and nature of synthesis errors in short-read sequencing, we analysed public datasets from 10X Genomics. Our initial assessment highlighted an elevated percentage of T bases in read1, corresponding to the barcode and UMI sequences. Particularly striking was the significant rise in T at the concluding base of the UMIs (**Fig. 1b, Supplementary Fig. 1, Supplementary Fig. 2, Supplementary Fig. 3**). This trend was even more pronounced in 10X Chromium Oxford Nanopore Technologies (ONT) long-read sequencing data (**Fig. 1c**). The increase in T base at the final position of the UMI potentially indicates the presence of bead truncation, suggestive of sequencing into the poly(dT) region. When examining Drop-seq^1^, we similarly observed an increase in the proportion of T throughout the UMI sequence in Illumina (**Fig. 1d**) but in ONT sequencing (**Fig. 1e**). Significant differences were observed in the spread of nucleotide bases along the lengths of UMIs across datasets. This implies that variations in estimating base molar weights before synthesis might influence the randomness of UMIs, thereby affecting their anticipated random distribution.

### Bead truncation results in diminished UMI complexity through T base overrepresentation

Having identified potential biases within both 10X Chromium and Drop-seq single-cell datasets, we next corroborated these findings through the sequencing of Drop-seq and 10X Chromium beads in isolation. This process revealed a predictable predominant peak size of 28bp for the 10X Chromium beads (**Fig. 2a**) and 20 bp for Drop-seq (**Fig. 2b**). Nonetheless, a notable truncation was observed, with merely 43.5% of the 10X Chromium beads and 35% of the Drop-seq beads exhibiting the anticipated length, demonstrating bead truncation. Subsequently, we analysed the counts for the top 20 UMIs in both 10X (**Supplementary Fig. 4**) and Drop-seq (**Supplementary Fig. 5**) experiments (**Fig. 2c, d**), uncovering a consistent trend of T base enrichment, especially at the end of UMIs. This further substantiates the occurrence of bead truncation.

**Figure 2:**
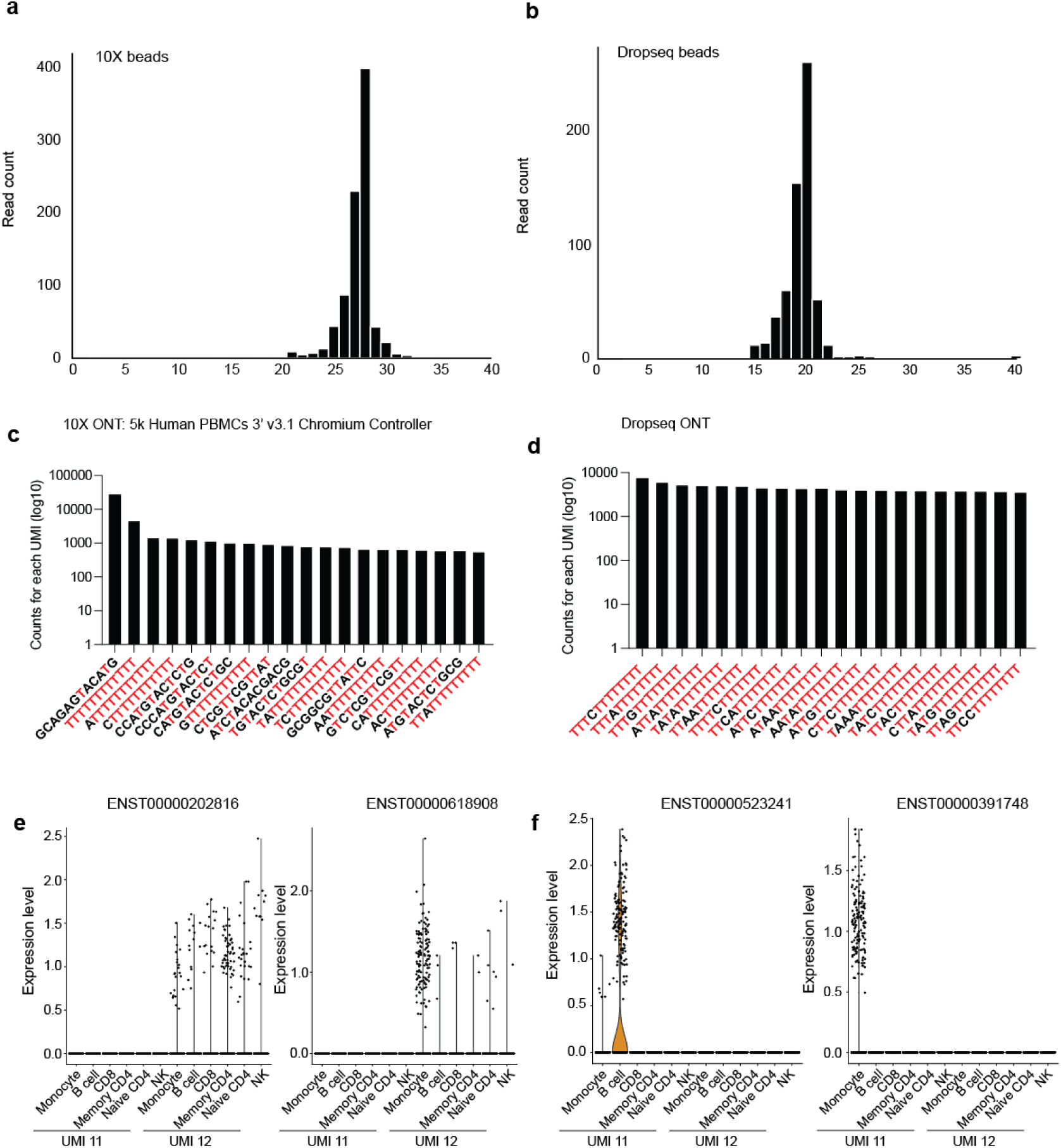
Impact of bead truncation on UMI deduplication accuracy. **a**, Generation of a sequencing library from 10X bead oligos and then sequenced using ONT flongle. **b**, A library was generated from oligos on Drop-seq beads and then sequenced using ONT flongle. **c**, Visualisation of the top 20 overrepresented UMIs in the 10X Chromium 5k PBMC v3.1 dataset, displaying the count of each UMI. **d**, Visualisation of the top 20 overrepresented UMIs in the Macosko et al 2015 dataset, displaying the number of UMI counts. **e**, Violin plots showing the expression of ENST00000202816 (ESF1) and ENST00000618908 (LAP3) across the cell annotated 10X Chromium 5k PBMC v3.1 dataset, analysed using UMI lengths of 11 or 12. **f**, Violin plots depicting the expression of ENST00000523241 (Paired Box 5; PAX5) and ENST00000391748 (Leukocyte immunoglobulin-like receptor subfamily B member 2; LILRB2) across the cells in the 10X Chromium 5k PBMC v3.1 dataset, offering significant into the significance of UMI length in gene expression analysis.

### Barcode truncation has minimal impact on cell identification

To elucidate the impact of potential synthesis inaccuracies on barcode assignment, we conducted a single-cell species mixing experiment using both the 10X Chromium and Drop-seq methodologies. For both methods, the barcode sequences were computationally truncated. Our results indicate that truncation does not adversely affect the quantification of cells identified (**Supplementary Fig. 6**). This outcome is attributed to the efficacy of the whitelisting correction strategy, which is capable of overcoming errors or truncations. Moreover, the barcode diversity is substantial for both 10X Chromium and Drop-seq, with each barcode separated by a minimum of at least 2 hamming distances from each other (**Supplementary Fig. 7**). The robustness of the whitelisting process against errors arising from synthesis, PCR, or sequencing was further validated by the computational shortening of barcodes, demonstrating no significant influence on the differentiation between human and mouse cells (**Supplementary Fig. 8**). This underscores the resilience and reliability of the whitelisting correction mechanism in maintaining accurate cell identification, despite potential synthesis errors.

### UMIs are significantly impacted by synthesis errors

Considering that UMI’s are intrinsically random, error correction cannot be facilitated through whitelisting. As a result, any error associated with UMIs may inadvertently lead to an overestimation of read counts and affect downstream differential expression. Our analysis identified notable increases in T bases at the end of UMI’s within 10X data (**Fig. 1**. and **Supplementary Fig. 1**), suggesting truncation and sequencing into the polyT region. Further examination into the effect of computationally truncating the UMI by a single base identified 115 differentially expressed transcripts between UMIs of 11 and 12 bases in length (**Supplementary Fig. 9 and Supplementary Table 1**).

It is noteworthy that no substantial enrichment of Gene Ontology (GO) terms was observed, implying that the differential expression of these genes might be attributable to stochastic variation rather than functionally coherence. Interestingly, the transcripts upregulated under the 12 base UMI condition showed a uniform increase across all cell types (**Fig. 2e**). In contrast, transcripts that show greater expressed in comparison to the 11-base UMI condition exhibited increases specific to certain cell types, aligning with their expected transcriptional activity (**Fig. 2f**). These findings suggest that truncation of UMI sequences can potentially compromise the accuracy of gene expression analyses. Our study underscores the critical point that while synthesis imperfections may not significantly affect barcode accuracy, they have implications for UMI identification, thereby impacting the fidelity of gene expression quantification.

### Inclusion of an anchor improves the ability to detect the start and end of the UMI

Having identified synthesis challenges associated with the bead oligonucleotides, we theorised that incorporating an anchor sequence between the barcode and UMI, and a V base between the UMI and the poly(dT) capture handle, could provide clearer demarcation of the beginning of the UMI. In our previous work^11^, we introduced the concept of a Common Molecular Identifier (CMI) sequence, designed to precisely quantify errors in a sequenced read (**Supplementary Fig. 10**). In the present study, we employed the CMI to evaluate the merits of incorporating an anchor within the oligonucleotide. To this end, we synthesised a free oligonucleotide that contained a PCR handle, a constant 12bp barcode region, preceded by a 4bp anchor, a 32 bp homodimer CMI, and finally a V base prior to the poly(dT) region (**Fig. 3a**). To pinpoint the CMI, we adopted two contrasting techniques. The first, known as the Positional Strategy, involved locating the end of the PCR handle through sequence alignment, then projecting the CMI’s onset to be 16 base pairs distant from this terminus. Conversely, the anchor strategy discerned the start of the CMI by pattern-matching the anchor sequence, thus identifying the CMI’s initiation immediately post the anchor position.

**Figure 3:**
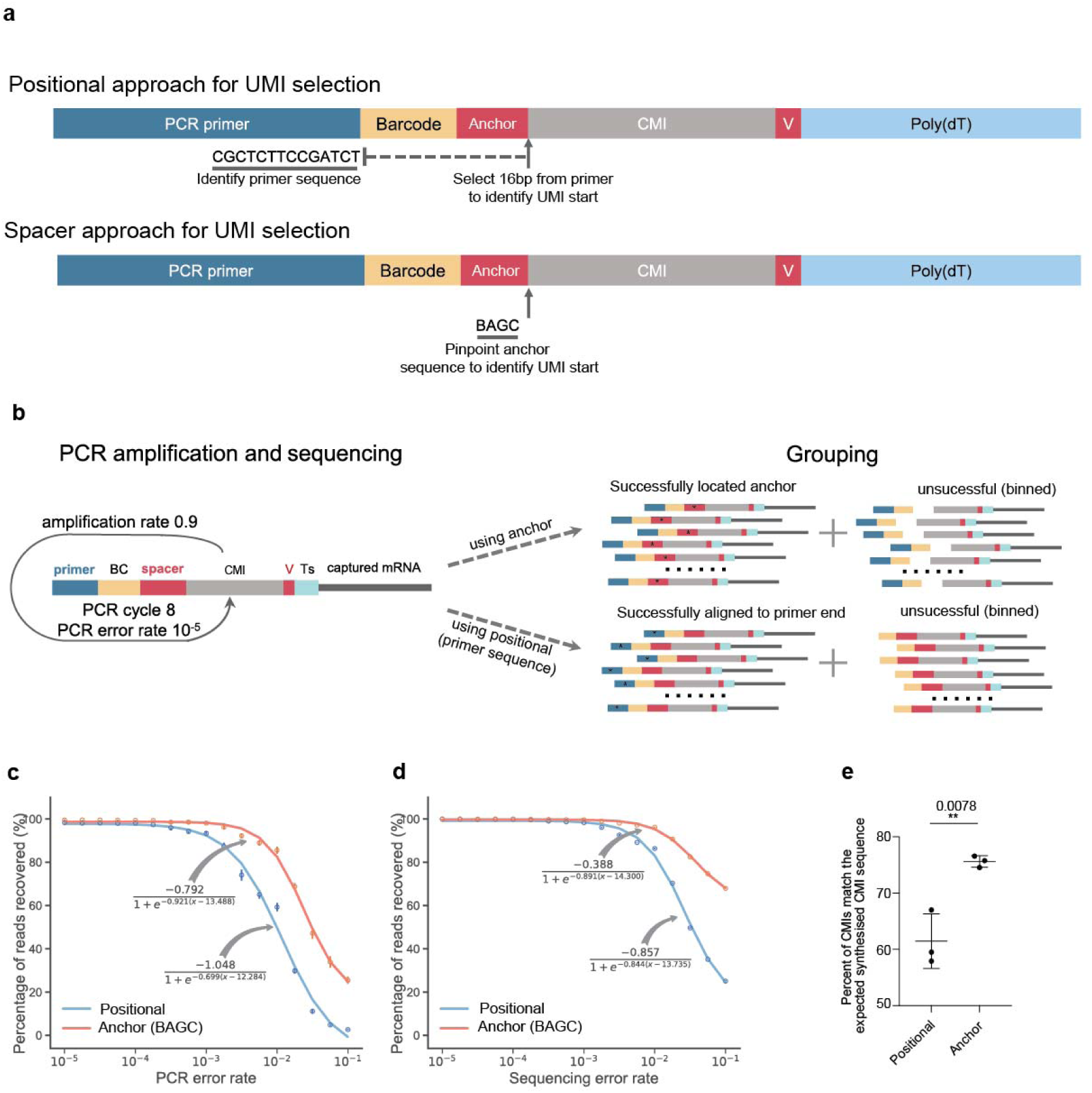
Inclusion of an anchor enhances the ability to identify the start of the CMI sequence. **a**. Schematic representation comparing two CMI selection methods: positional and anchor. The positional method identifies the PCR primer’s end through local alignment and selects a site 16 base pairs downstream. In contrast, the anchor method utilizes a regular expression to pinpoint the CMI’s start position. **b**. A schematic showing the workflow of bead simulation for evaluating the UMI identification using the anchor and positional strategies. **c,** The number of reads recovered were plotted as a function of the PCR error rate**. d,** The number of reads recovered were plotted as a function of the sequencing error rate. **e**. A bulk reverse transcription and PCR was performed using the oligonucleotide design in **a**. The percentage of CMIs that perfectly matched the anticipated CMI sequence were measured for both the positional and anchor strategy. **e**, Data analysed using Unpaired t-test assuming both populations have the same standard deviation. ** P<0.01. The mean and S.D. are plotted.

Based on the specified design of the bead, an initial computational analysis was undertaken to assess the efficiency of capturing CMIs, using simulated data (**Fig. 3b, c**). This analysis, supported by benchmarking simulations, provides insights into the potential for enhancing CMI identification through two distinct methodologies. Initially both methodologies demonstrate comparable effectiveness in identifying CMIs. However, as the PCR and sequencing error rates increased, the anchor strategy significantly outperformed the positional strategy. To offer a more universal comparison of the efficiencies of these methods across various scenarios, logistic functions were employed. (**Fig. 3c,d and Supplementary Fig. 11**). Next, we experimentally evaluated the two strategies using an mRNA bulk library preparation experiment, followed by ONT sequencing. We observed a significant increase in the accurate identification of CMIs using the anchor method compared to the positional one (**Fig. 3d**). This data indicates that incorporating an anchor can boost the precision of determining the CMI’s initial position.

### New bead design containing an anchor mitigates oligo truncation and coupling errors

After demonstrating that incorporating an anchor between the barcode and UMI enhances UMI identification, we integrated this feature into our Drop-seq bead designs. Previously, we developed an advanced Drop-seq method named scCOLOR-seq^12, 13^, which employs error-correcting homodimer UMIs. By adding both an anchor and a V base to the capture oligonucleotide, we reinforced these beads to ensure a clear separation between different components, which paves the way for more accurate UMI recognition (**Fig. 4a**). The inclusion of an anchor clearly demarked the boundary between the barcode and the UMI sequence (**Fig. 4b**). Our primary objective was to assess whether this modification increased the accuracy of UMI sequence detection.

**Figure 4:**
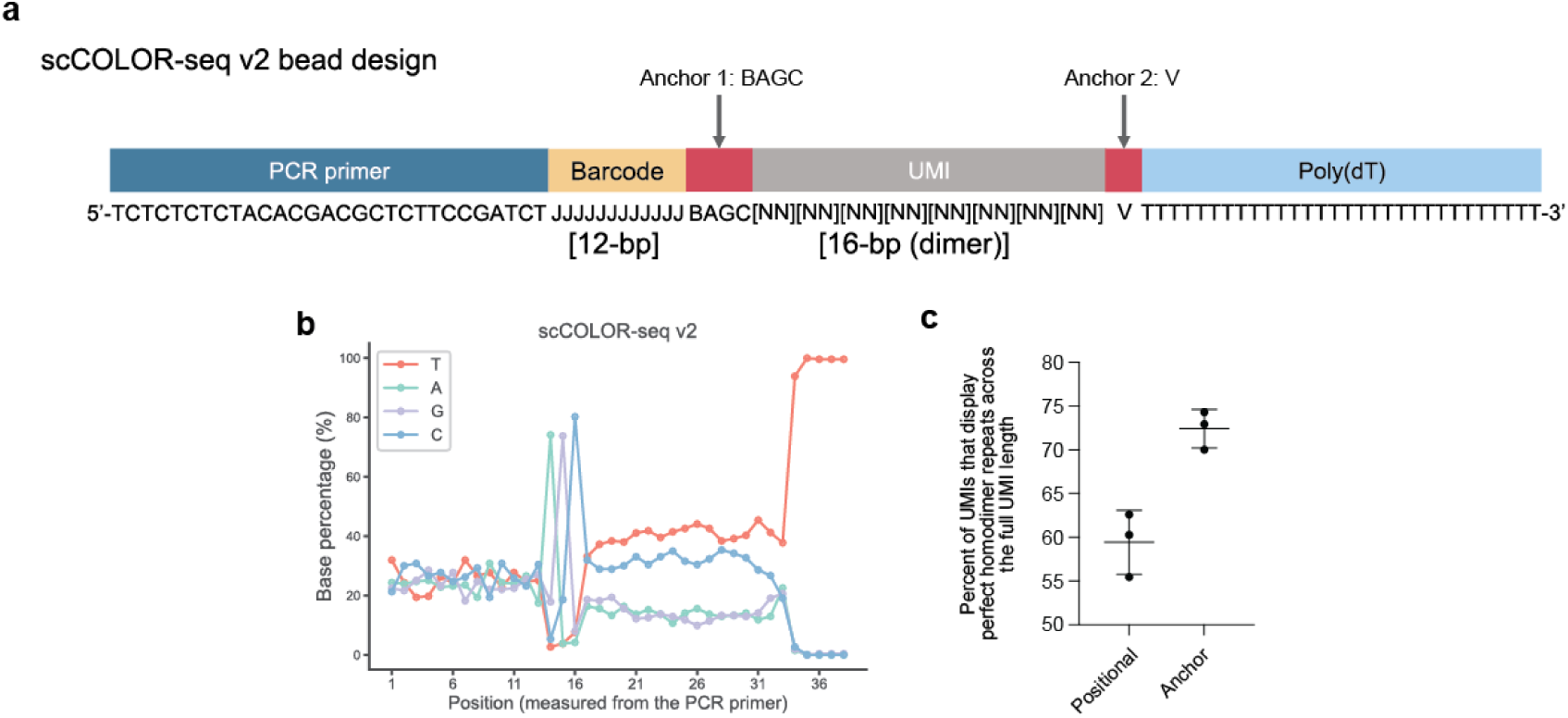
The inclusion of an anchor in the mRNA capture beads improves the UMI recovery. **a,** A schematic showing the scCOLOR-seq v2 bead design that incorporates an anchor sequence between the barcode and UMI, in addition to a homodimer UMI followed by a V base between the poly(dT) capture region. **b**, The percent of A, C, G and T base plotted for each base within the fastq file upstream of the PCR primer end position. **c**, The homodimer repeats were used as a proxy for measuring improved UMI selection. The percent of perfect homodimer repeats across the full length of the UMI is plotted for using the positional and the anchor approach.

Given the UMI’s unique homodimer composition, we theorised that assessing perfect dimer nucleotide concordance throughout the UMI would be an effective validation metric for our improved strategy. If the homodimer concordance improved, it would suggest better UMI identification. We next compared the positional and anchor approaches. The addition of the anchor indeed increased the homodimer concordance rate (**Fig. 4c and Supplementary Fig. 12**), underscoring our ability to precisely determine the UMI’s boundaries. This marked rise in the dimer concordance rate for the homodimer UMI bolstered our initial theory, highlighting that the anchor’s inclusion augments the UMI sequence identification process.

### The inclusion of an anchor improves UMI counts and number of transcripts detected

We next tested the new bead design in a species mixing experiment. During the subsequent data analysis, we compared both the positional and anchor techniques. Notably, implementing the anchor-centric method resulted in a significant increase in the number of detected UMIs (**Fig. 5a**) and transcripts (**Fig. 5b**) per cell relative to the positional strategy. Furthermore, greater number of features were identified within both human (**Fig. 5c**) and mouse (**Fig. 5d**, **Fig. 5e and Fig. 5f**) cells using the anchor when compared to the positional approach. This suggests that integrating an anchor in the bead oligonucleotide sequence effectively minimises artefacts, yielding a more precise tally of unique molecules across a broad spectrum of features.

**Figure 5:**
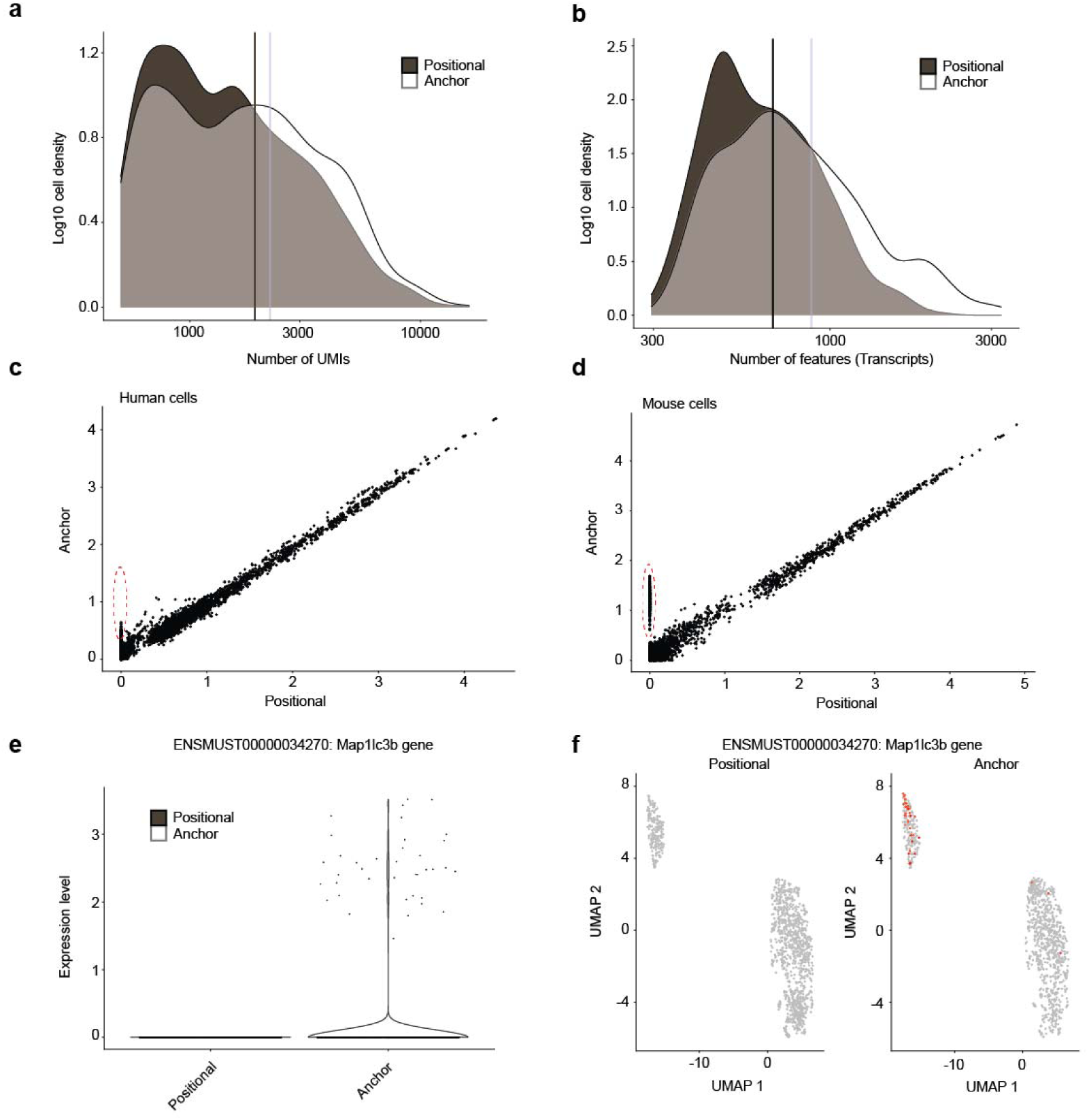
Including an anchor into the beads increases the number of UMIs and transcripts detected. **a**, The log10 density of UMIs per cell comparing the positional and anchor UMI selection methods. **b**, The log10 density of transcripts per cell for both positional and anchor UMI selection approaches. **c**, Correlation of transcripts in human-origin cells between the positional and anchor approaches. Transcripts with enhanced detection using the anchor method are highlighted with a red cicle. **d**, Correlation of mouse-origin transcripts between the positional and anchor approach, with the red circle identifying transcripts more readily detected using the anchor UMI identification approach. **e**, Expression profile of the transcript ENSMUST00000034270 for the positional and anchor approach. **f**, UMAP plots showing the increased expression of ENSMUST00000034270 using the anchor approach. For plots **c**-**f**, each dot represents a single cell. Data shown is from one of three independent experiments.

## Discussion

Single-cell RNA-sequencing (scRNA-seq) technology is a rapidly advancing technology that is revealing new scientific findings^14, 15^. However, the data it generates can often contain numerous technical biases^16^. In this study, we evaluated bead synthesis errors in droplet-based single-cell sequencing. We identified several technical inaccuracies arising from imperfect synthesis of bead-bound oligonucleotides. To address these errors, we devised an enhanced bead design, which improved barcode and CMI assignment. One of the pressing issues with scRNA-seq data is the amplification of technical errors, which introduces noise^17, 18^, making gene counting unreliable and potentially skewing differential gene expression results. The ideal scenario would see scRNA-seq tools accurately gauging and correcting uncertainties from these biases and errors^19, 20^. However, discerning technical errors from genuine biological variations using computational techniques alone is challenging, often because of a lack of ground truth in experimental data. In earlier studies we have identified sequencing and PCR errors as sources of technical inconsistencies in single-cell transcriptomics^11, 12^, causing inaccurate feature counting. Yet, the impact of oligonucleotide bead synthesis on droplet-based single-cell sequencing has been largely unexplored.

There are two primary bead synthesis strategies: on-bead chemical synthesis and enzymatic ligation. The on-bead chemical method incorporates a barcode generated using a split and pool technique, and then a random UMI synthesised using single random nucleosides, a concept first introduced by Drop-seq^1^. Its efficient creation of capture oligonucleotides on beads offers a diverse barcode size and ensures most cells align with a bead. Conversely, the enzymatic ligation method, initially introduced by InDrops^2^ and adopted by other techniques^3, 21^, presents a more modular approach to bead synthesis. In this strategy, bead fabrication utilises combinations of a limited set of pre-synthesised oligonucleotides. In alignment with the chemical synthesis method’s principles, the barcode is created using a split and pool approach using small stretches of pre-synthesised building blocks. However, a distinct feature here is that a pre-synthesised random UMI is enzymatically ligated to the barcode. While these methods differ in their approach to bead synthesis, both exhibit a notable drawback. As we show that both are prone to significant truncation, likely due to challenges in purifying the oligonucleotides once affixed to the beads. The immobilised oligonucleotides can’t be cleaved and purified due to the cell barcodes contained within each one.

We have shown that synthesis issues significantly influence the precision of single-cell sequencing. Oligonucleotide synthesis errors, with an average rate of 1 in 100 bases^10, 22^, commonly result in truncated or elongated oligonucleotides. While standard oligonucleotide synthesis allows for the separation and purification of truncated sequences from full-length products using HPLC or PAGE, these methods are not feasible for single-cell sequencing applications. As a result, these length discrepancies can interfere with barcode and UMI sequence detection, a problem further amplified in long-read sequencing. This is because, in long-read sequencing, barcodes are discerned following computational alignment to the PCR primer, coupled with positional matching to pinpoint the cell barcode and UMI’s beginning^23, 24^. Variability in oligonucleotide lengths on beads makes barcode and UMI detection more error prone. Our findings suggest that identifying the barcode and UMI can be challenging for both Drop-seq and 10X beads because of synthesis errors. UMIs present a larger issue than barcodes; while whitelisting can correct most barcode errors, the UMIs’ inherent randomness makes whitelisting impossible. To tackle synthesis-related challenges with UMI detection, we incorporated an anchor and synthesised our UMIs using homodimer nucleoside amidites. This strategy not only pinpoints the start of the UMI but also identifies and fixes issues in the UMI arising from PCR, sequencing, and synthesis. As the field of single-cell biology advances and its techniques find application in clinical settings^14, 25, 26^, the development of more rigorous methods to ensure the reliability of single-cell methodologies becomes imperative. Our findings demonstrate that the incorporation of spacers mitigates synthesis-related errors, thereby enhancing the accuracy of differential expression analysis.

## Methods

### Cell lines and reagents

Jurkat cells were purchased from ATCC. 5TGM1 were a kind gift from Prof Clair Edwards. Both Jurkat and 5TGM1 cells were cultured in complete RPMI medium supplemented with 10% Foetal Calf Serum. All parental cell lines were tested twice per year for mycoplasma contamination and authenticated by STR during this project.

### Oligonucleotide synthesis

Single-cell oligonucleotide bead synthesis was executed in line with previously established methods^12, 13^, but with the following alterations to the bead design:

5’-Bead-HEG_Linker-TCTCTCTCTACACGACGCTCTTCCGATCTJJJJJJJJJJJJBAGCNNNNNNNNVTTTTTTTTTTTTTTTTTTTTT TTTTTTTTT-3’

Here, "J" represents the monomer split-and-pool barcode, while "N" signifies the dimer amidite UMI and B signifies either a C, G or T. We procured CMI oligos from Sigma-Aldrich with desalting, designed as follows:

Anchor oligo: 5’-TCTCTCTCTACACGACGCTCTTCCGATCTAGTGCGTAGCTGBAGCGGAACCTTGGCCTTAATTGGTTAA GGTTGGAATTTTTTTTTTTTTTTT-3’

No anchor: 5’-TCTCTCTCTACACGACGCTCTTCCGATCTAGTGCGTAGCTGGGAACCTTGGCCTTAATTGGTTAAGGTTGGAATTTTTTTTTTTTTTTT-3’

### Bead sequencing library preparation

2,000 beads (in TE/TW storage buffer) were added to a PCR tube. Beads were washed in 200 ul of H2O, centrifuged (100 g, 1 minute) and supernatant removed. 50 ul of PCR mix (1.5 ul indexed polyA primer (100 uM), 1.5 ul new P5 primer (100 uM; for beads carrying a SMART PCR handle) or 1.5 ul NEBNext i50x primer (100 uM; for beads carrying P5 PCR handle) or 10 ul SI primer (from 10X kit; for 10X beads), 25 ul KAPA HiFi master mix and H2O to 50 ul) was added, mixed and immediately run in a thermocycler with the following conditions. 95°C for 3 minutes. 12 cycles of: 95°C for 20 seconds, 60°C for 15 seconds, 72°C for 15 second then 72°C for 1 minute and 4C hold. After PCR, samples were cleaned up by adding 100 ul of SPRIselect beads (Beckman Coulter) and following the manufacturer’s instructions. Samples were eluted in 20 ul of H2O and run on a HS D1000 tape (Agilent). ONT Flongle libraries were prepared and loaded according to protocol sqk-lsk114-ACDE_9163_v114_revJ_29Jun2022-flongle, starting with 100 fmol of PCR product (or pooled products where indexing was used). 10 fmol of library was loaded onto a flongle flow cell. Sequencing was run for 24 hours using super-accuracy basecalling, minimum fragment size of 20 bp and no filtering based on Q score. Primers used for the PCR and sequencing:

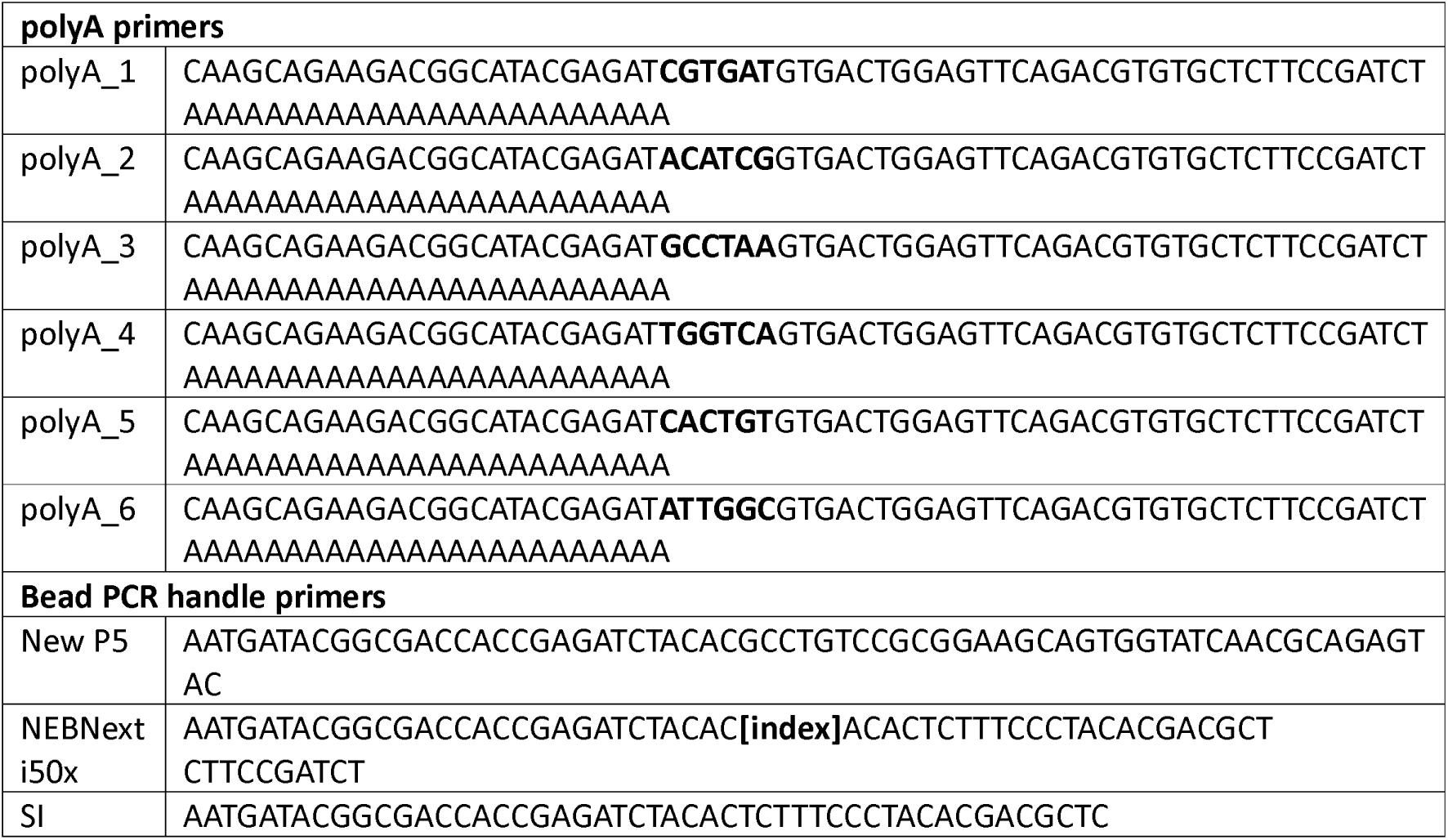

### 10X chromium library preparation

We prepared a single-cell suspension using Jurket and 5TGM1 cells using the standard 10X Genomics chromium protocol as per the manufacturer’s instructions. Briefly, cells were filtered into a single-cell suspension using a 40 µM Flomi cell strainer before being counted. We performed 10X Chromium library preparation following the manufacturers protocol. Briefly, we loaded 3,300 Jurkat:5TGM1 cells at a 70:30 split into a single channel of the 10X Chromium instrument. Cells were barcoded and reverse transcribed into cDNA using the Chromium Single Cell 3’ library kit and get bead v3.1. We performed 10 cycles of PCR amplification before cleaning up the library using 0.6X SPRI Select beads. The library was split and a further 20 or 25 PCR cycles were performed using a biotin oligonucleotide (5-PCBioCTACACGACGCTCTTCCGATCT) and then cDNA was enriched using DynabeadsTM MyOneTM streptavidin T1 magnetic beads (Invitrogen). The beads were washed in 2X binding buffer (10mM Tric-HCL ph7.5, 1mM EDTA and 2M NaCl) then samples were added to an equi-volume amount of 2X binding buffer and incubated at room temperature for 10 mins. Beads were placed in a magnetic rack and then washed with twice with 1X binding buffer. The beads were resuspended in H2O and incubated at room temperature and subjected to long-wave UV light (∼366 nm) for 10 minutes. Magnetic beads were removed, and library was quantified using the QubitTM High sensitivity kit. Libraries were then prepared before sequencing.

### Drop-seq and scCOLOR-seqv2 library preparation

Single-cell capture and reverse transcription were performed as previously described (ref). Briefly, Jurkat and 5TGM1 cells (20:80 ratio) were filtered into a single-cell suspension using a 40 µM Flomi cell strainer before being counted. Cells were loaded into the DolomiteBio Nadia Innovate system at a concentration of 310 cells per µL. Custom synthesised beads were loaded into the microfluidic cartridge at a concentration of 620,000 beads per mL. Cell capture was then performed using the standard Nadia Innovate protocol according to manufacturer’s instructions. The droplet emulsion was then incubated for 10 mins before being disrupted with 1H,1H,2H,2H-perfluoro-1-octanol (Sigma) and beads were released into aqueous solution. After several washes, the beads were subjected to reverse transcription. Prior to PCR amplification, beads were treated with ExoI exonuclease for 45 min. PCR amplification was then performed using the SMART PCR primer (AAGCAGTGGTATCAACGCAGAGT) and cDNA was subsequently purified using AMPure beads (Beckman Coulter). The library was split and a further 20 or 25 PCR cycles were performed using a biotin oligonucleotide (5—PCBioTACACGACGCTCTTCCGATCT) and then cDNA was enriched using DynabeadsTM MyOneTM streptavidin T1 magnetic beads (Invitrogen). The beads were washed in 2X binding buffer (10mM Tric-HCL ph7.5, 1mM EDTA and 2M NaCl) then samples were added to an equi-volume amount of 2X binding buffer and incubated at room temperature for 10 mins. Beads were placed in a magnetic rack and then washed with twice with 1X binding buffer. The beads were resuspended in H2O and incubated at room temperature and subjected to long-wave UV light (∼366 nm) for 10 minutes. Magnetic beads were removed, and library was quantified using the QubitTM High sensitivity kit. Libraries were then prepared for sequencing.

### Single-cell Nanopore library preparation for sequencing

A total of 500 ng of single-cell PCR input was used as a template for ONT library preparation. Library preparation was performed using the SQK-LSK114 (kit V14) ligation sequencing kit, following the manufacturers protocol. Samples were then sequenced on either a Flongle^TM^ device or a PromethION^TM^ device using R10.4 (FLO-PRO114M) flow cells. The fast5 sequencing data was basecalled to fastq files using guppy basecaller (v6.5.7) using the Super-accuracy mode within the MinKnow software (v23.04.6).

### Read simulation

To understand how much improvement in successfully localising the start of UMIs can be gained from our optimized bead design, we conducted a series of in silico experiments by simulating beads at different application scenarios, including PCR/sequencing substitution errors and PCR/sequencing indels (i.e., insertion and deletion errors). Due to the known composition of the proposed bead, we began directly by amplifying one bead at a single genomic locus over a predefined number of PCR cycles. The overall parameters for simulation were set as follows. Reads were PCR amplified over 8 cycles with an amplification efficiency rate of 0.9. Both PCR and sequencing error rates varied from *10* ^-5^ to *10* ^-1^. The PCR and sequencing error rates were fixed to be *10* ^-5^ and *10* ^-3^, respectively, when another parameter needed to vary. As suggested by Potapov et al^27^, we set PCR deletion and insertion rates as 2.4 x 10^-6^ and 7.1 x 10^-7^, respectively, but varied them from *10* ^-5^ to *10* ^-1^, so did sequencing deletion and insertion rates. After PCR amplification, 5000 reads were subsampled for sequencing. The numbers of both substitution errors and indels were determined from negative binomial distributions. The positions of the error bases were randomly picked. To minimise the variance, we executed 10 permutation tests for each application scenario.

### Simulation for comparing positional and anchor schemes

To precisely localise the UMI from a read, our proposed anchor strategy involved incorporating an anchor interposed between a barcode and a UMI as well as a V base placed after the UMI for demarcating its followed poly(dT)s. To ascertain the improvement of identifying UMIs by this design, we further implemented a positional scheme for comparison, which supposedly reaches the entrance to the UMI by counting 16 bases from an ending string of 14 bases in the primer. To test the robustness of the two strategies in coping with high error rates, we subjected simulated reads into either PCR amplification or sequencing at error rates of *10* ^-5^, 2.5 x 10^-5^, 5x 10^-5^, 7.5 x 10^-5^, 0.0001, 0.00025, 0.0005, 0.00075, 0.001, 0.0025, 0.005, 0.0075, 0.01, 0.025, 0.05, 0.075, and 0.1. To evaluate their efficiency, we calculated P, defined as

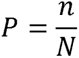

where *N* is the subsampled 5000 reads and n represents the number of reads whose UMIs can successfully be discovered by using the anchor or positional strategy.

### Simulation for UMI identification with or without an anchor

It has been widely considered to be difficult to correctly extract UMIs from reads that suffer indels. The inclusion of an anchor in between a UMI and a segment of poly(dT)s makes it possible to be protected from being contaminated by reading Ts into the UMI portion, while the inclusion of an anchor in between a cell barcode and a UMI can set a buffer that suffices to separate the UMI from the cell barcode, which leaves room for UMI identification. To quantify the influence of indels on reads, we simulated scCOLOR-seq reads by adding indels during the PCR amplification or sequencing stage at error rates of *10* ^-5^, 2.5 x 10^-5^, 5x 10^-5^, 7.5 x 10^-5^, 0.0001, 0.00025, 0.0005, 0.00075, 0.001, 0.0025, 0.005, 0.0075, 0.01, 0.025, 0.05, 0.075, and 0.1. To evaluate their efficiency, we constructed a Q value, computed by

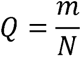

where *m* represents the number of reads whose UMIs can successfully be discovered if the anchor (BAGC) is used, and the number of reads that are free from indels (taken as a strategy for evaluation without an anchor), otherwise.

### Characterization of the quantity of reads captured by successfully identifying their UMIs

We show that a logistic function can be used to characterize and describe how the number of successfully captured reads (i.e., *P* and *Q*) varies against both substitution errors or indels. The logistic function *l* has the following form.

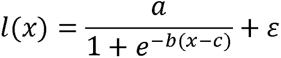

where *a*, b, c, and *ε* are four parameters to be estimated according to the computed *P* and *Q* values. Each logistic equation reflects the influence of substitution errors or indels imposed on reads captured by successfully identifying their UMIs and represents the robustness of our proposed and compared methods in UMI identification.

### Edit distance-based similarity of barcodes

To infer the extent to which any two barcodes in the whitelist can change to one another when they suffer from errors, we made an all-against-all comparison between barcodes in the chemistry V2 and V3 whitelists, which were downloaded from https://teichlab.github.io/scg_lib_structs/methods_html/10xChromium3.html. There are 737,280 barcodes for chemistry V2 and 6,794,880 barcodes for chemistry V3. To accelerate the calculation, we employed a distributed computing strategy (**Supplementary Figure 7a**). To illustrate our approach, we detail the comparison of barcodes utilising the V3 chemistry against a predefined whitelist. Our computational framework initiated 680 discrete tasks for execution on our server cluster. Within each task, edit distances were computed for 10,000 selected barcodes against those previously unexamined. Following a complete comparative analysis, a barcode was eliminated from the dynamically updated whitelist, precluding further comparisons. This method facilitated the execution of comprehensive pairwise comparisons among barcodes (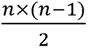, where *n* represents the total count of barcodes within the whitelist), thereby enhancing computational efficiency. The outcomes of these comparisons were systematically documented in JSON format upon the conclusion of each barcode’s edit distance evaluation. Subsequent to the completion of all tasks, these JSON records were amalgamated into a singular comprehensive file.

### 10X Genomics datasets of short-reads

To comprehensively understand how genomic positions from sequencing reads are represented by different nucleotides, we conducted a large-scale analysis over 41 scRNA-seq datasets downloaded from 10x Genomics using chemistry v2, v3, and v3.1 (**Supplementary Table 2, Supplementary Table 3 and Supplementary Table 4**). To make the data representative of universality and diversity, we chose these datasets with a few criteria, such as from a variety of species, including human, mouse, and their mixture, and from cells or nuclei.

### 10X chromium short-read analysis workflow

The data was processed using a custom CGAT-core^28^ pipeline ‘pipeline_10x_shortread’, which is included within the TallyTriN Github repository (https://github.com/cribbslab/TallyTriN/blob/main/tallytrin/pipeline_10x_shortread.py). Briefly, the quality of each fastq file is evaluated using fastqc toolkit (ref) and summary statistics collated using Multiqc (ref). We then identify putative barcodes using UMI-tools whitelist module and then extract the barcodes and UMIs from the read 1 fastq file and append them onto the read 2 file using umi_tools extract module. Hisat2 (ref) is then used to map the reads to the hg38_ensembl98 genome and the resulting bam file is then sorted, and each read is assigned to a feature using featureCounts (ref), with the alignment written to the XT flag of the output bam file. This is then indexed using samtools (ref) and then UMI counting is performed using the umi_tools count module before being converted to a market matrix format. The resulting matrix files are then parsed into R/Bioconductor (v4.0.3) using the BUSpaRse (v1.14.1) package and downstream analysis was performed using Seurat (v 4.3.0.1). Transcript matrices were cell-level scaled and log-transformed. The top 2000 highly variable genes were then selected based on variance stabilising transformation which was used for principal component analysis (PCA). Clustering was performed within Seurat using the Louvain algorithm. To visualise the single-cell data, we projected data onto a Uniform Manifold Approximation and Projection (UMAP).

### 10X chromium long-read analysis workflow

To analyse the 10X chromium long-read data, we developed a custom cgatcore pipeline named ‘pipeline_10x’ in the TallyTriN Github repository (https://github.com/cribbslab/TallyTriN/blob/main/tallytrin/pipeline_10x.py). We split the fastq file into segments to optimize processing time. Each read’s polyA tail was identified and reverse complemented for consistent orientation, discarding reads without a polyA tail. We identified the barcode and UMI by locating the PCR primer sequence ‘AGATCGGAAGAGCGT’ through pairwise alignment, and then extracted the 16 bp barcode and 12 bp UMI based on position. Barcodes were corrected using a whitelisting method similar to UMI-tools.

Mapping was done using minimap2 (v2.22) with settings: -ax splice -uf MD –sam-hit-only – junc-bed, referencing the human hg38 and mouse mm10 transcriptomes. The resultant sam file was arranged and indexed via samtools. Read counting employed UMI-tools’ count module, converting counts to a matrix format. We processed the raw expression matrices using R/Bioconductor (v4.0.3) and devised scripts to depict barnyard plots, displaying mouse and human cell proportions. Matrices were cell-level scaled and centre log ratio transformed. We selected the top 2000 variably expressed genes post-variance stabilising transformation for PCA. Clusters were identified in Seurat using the Louvain algorithm. For visualisation, we projected the single-cell data onto a Uniform Manifold Approximation and Projection (UMAP).

### Drop-seq analysis workflow

Drop-seq data was processed using a CGAT-core workflow ‘pipeline_macosko’, which is included within the TallyTriN repository (https://github.com/cribbslab/TallyTriN/blob/main/tallytrin/pipeline_singlecell_macosko.py) . Briefly, the fastq file was split into chunks so that analysis scripts can be processed on sections of the data to reduce the processing time. The polyA tail for each read was identified and then reverse complemented to keep all the reads within the same orientation, any reads not containing a polyA tail were discarded. Next the barcode and UMI was identified based on the identification of the PCR primer sequence using pairwise alignment and then selecting the 12 bp barcode and the 8bp UMI using positional matching. Next, the barcodes were corrected using a whitelisting approach like the one implemented by UMI-tools. The reads were then merged and then mapping was performed using minimap2 (v2.22) (ref). Mapping settings we as follows: -ax splice -uf MD –sam-hit-only –junc-bed and using the reference transcriptome for human hg38 and mouse mm10. The resulting sam file was sorted and indexed using samtools. Mapping settings we as follows: -ax splice -uf MD – sam-hit-only –junc-bed and using the reference transcriptome for human hg38 and mouse mm10. The resulting sam file was sorted and indexed using samtools. Counting was performed using UMI-tools count module before being converted to a market matrix format. Raw transcript expression matrices generated were processed using R/Bioconductor (v4.0.3) and custom scripts were used to generate barnyard plots showing the proportion of mouse and human cells. Transcript matrices were cell-level scaled and centre log ratio transformed. The top 2000 highly variable genes were then selected based on variance stabilising transformation which was used for principal component analysis (PCA). Clustering was performed within Seurat using the Louvain algorithm. To visualise the single-cell data, we projected data onto a Uniform Manifold Approximation and Projection (UMAP).

### scCOLOR-seqv2 analysis workflow

To process the drop-seq data, we wrote a custom cgatcore pipeline (https://github.com/cribbslab/TallyTriN) (ref). We followed the workflow previously described for identifying barcodes and UMIs using scCOLOR-seq sequencing analysis (ref). Briefly, to determine the orientation of our reads, we first searched for the presence of a polyA sequence or a polyT sequence. In cases were the polyT was identified, we reverse complemented the read. We next identified the barcode sequence by searching for the polyA region and flanking regions before and after the barcode. The dimer UMI was identified based upon the primer sequence TCTTCCGATCT at the TSO distal end of the read. Barcodes and UMIs that had a length less than 50 base pairs were discarded. Next, the barcodes were corrected using a whitelisting approach like the one implemented by UMI-tools. The reads were then merged and then mapping was performed using minimap2 (v2.22) (ref). Mapping settings we as follows: -ax splice -uf MD –sam-hit-only –junc-bed and using the reference transcriptome for human hg38 and mouse mm10. The resulting sam file was sorted and indexed using samtools. Mapping settings we as follows: -ax splice -uf MD – sam-hit-only –junc-bed and using the reference transcriptome for human hg38 and mouse mm10. The resulting sam file was sorted and indexed using samtools. The transcript name was then appended to the bam XT flag using the xt_tag_nano script before umi_tools count module was used to count the features using the following settings: --per-gene --gene-tag=XT --per-cell --dual-nucleotide. The umi_tools used to correct for the dimer UMIs is located in the AC-dualoligo in a fork at the repository: https://github.com/Acribbs/UMI-tools. Downstream analysis in R was then performed as described for the Drop-seq analysis workflow above.

### Data availability

Source data is provided with this manuscript. All custom pipelines used within this analysis are available on github (https://github.com/cribbslab/TallyTriN). Sequencing data have been deposited in the GEO under accession number GSE263458.

## Author contributions

A.P.C. designed the study with contributions from M.P., T.B. Jr, T.B. Sr and U.O. A.P.C. and J.S conducted data analysis, generated the figures and wrote the paper with input from all authors. J.S. conducted data analysis, generated figures and wrote the paper. M.P., D.L., S.H., G.H., J.R., N.M., E.C. and V.G. performed experiments.

