## Supplementary Figures for "Anchor-Enhanced Bead Design for Reduced Oligonucleotide Synthesis Errors in Single-cell sequencing"

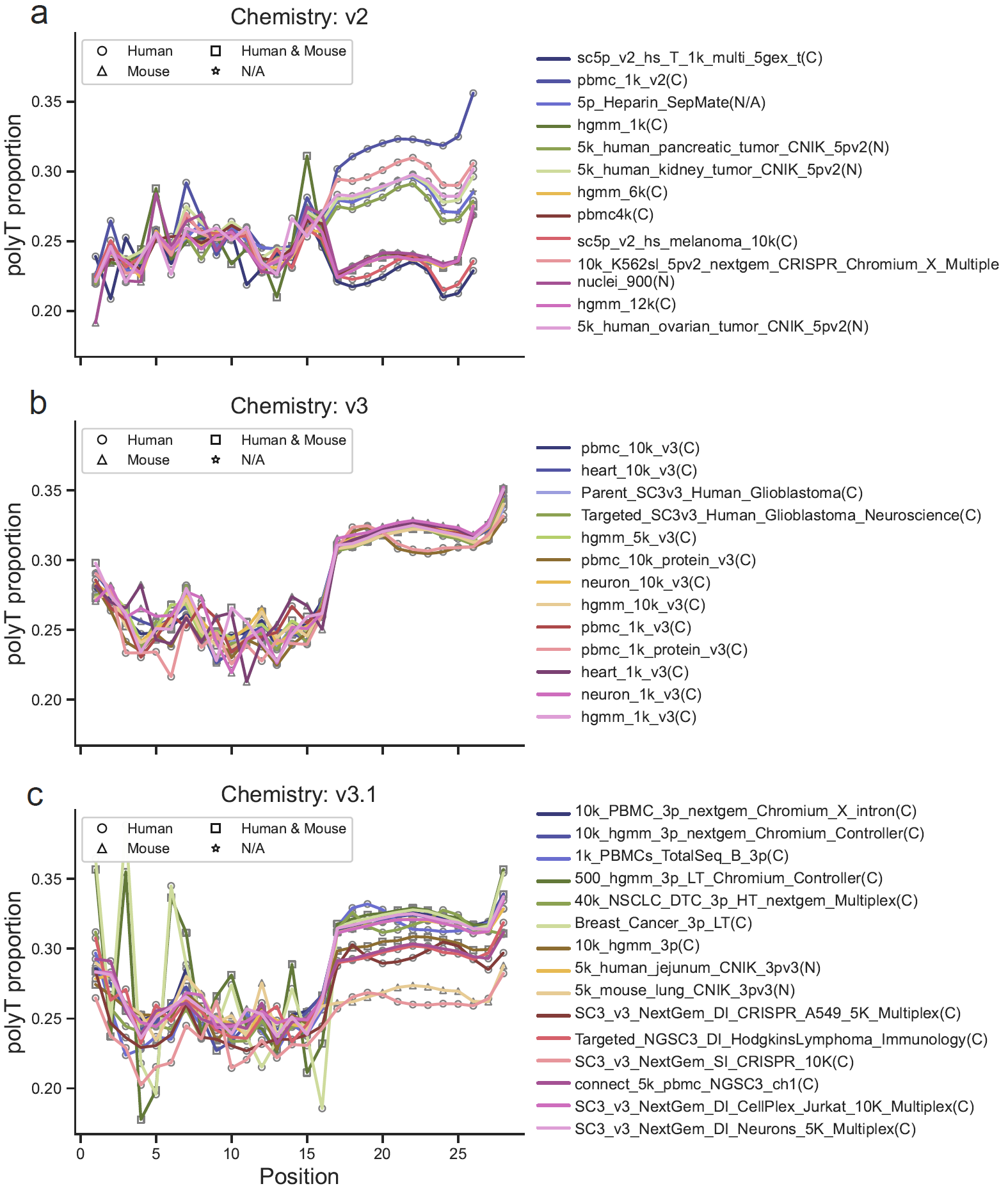

**Supplementary Figure 1:** Comparative Analysis of Thymine (T) Base Proportion in 10X Beads Across Versions. This figure delineates the frequency distribution of thymine (T) bases in two different versions of 10X beads. **a**) illustrates the prevalence of T bases in version 2 of the 10X beads, using data obtained from public datasets. **b**) represents the frequency of T bases in version 3 of the 10X beads. **c**) represents the frequency of T bases in version 3.1 (updates for version 3) of the 10X beads. This comparison includes a total of 43 datasets (**Supplementary Table X** for details) and aims to elucidate potential variations in base composition between the two types of bead versions. C: sampled from cells. N: sampled from nuclei. N/A: not available.

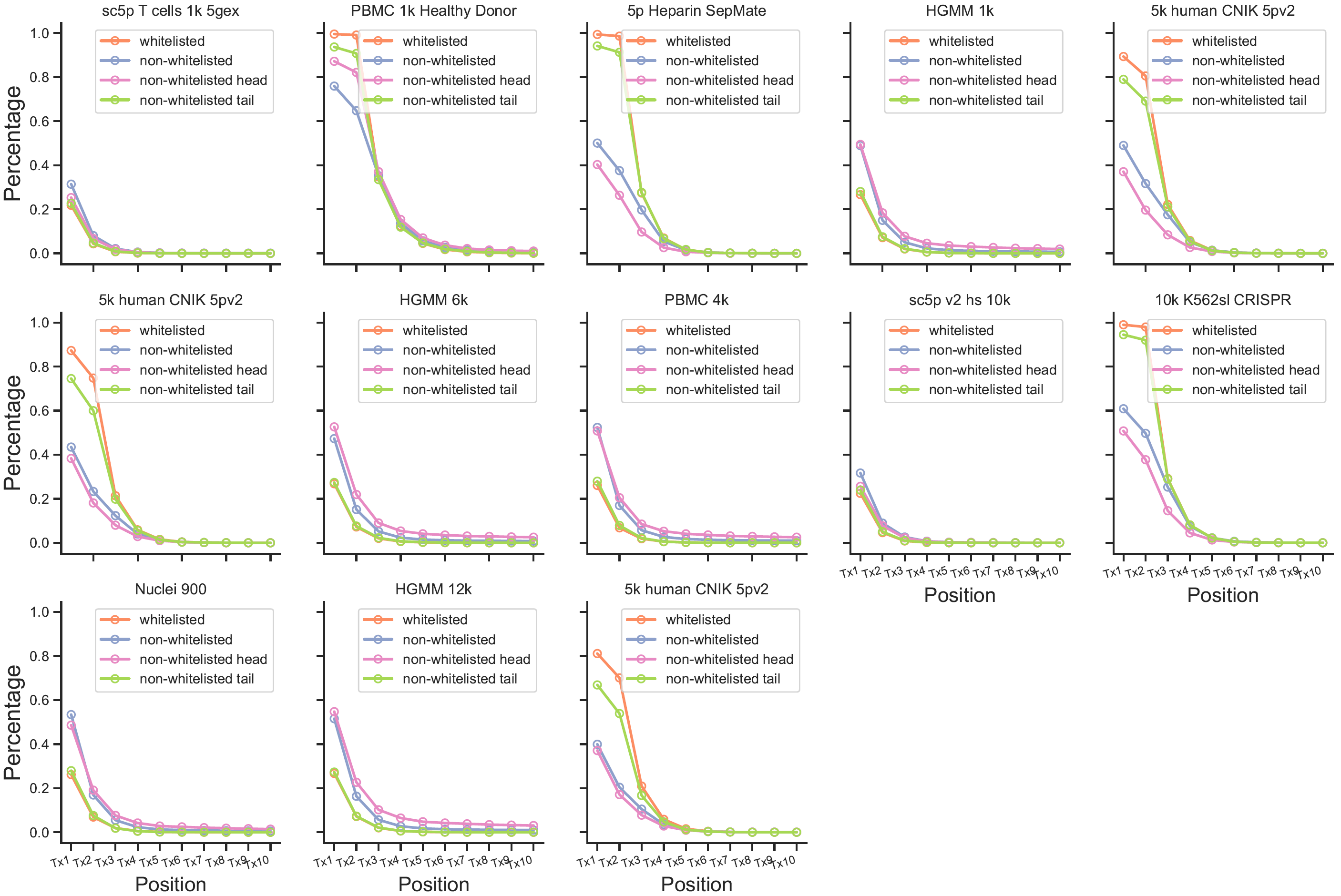

**Supplementary Figure 2**. Percentage of Ts in UMIs across 13 datasets using chemistry V2 with respect to their barcodes that are or are not whitelisted or those non-whitelisted barcodes that have their head/tail regions overlapped with head/tail regions of barcodes in the whitelist. Each of the barcodes is split into two parts with the first 8 nucleotides denoted as the head region and the last 8 nucleotides denoted as the tail region. We observe the percentage of T increases towards the end of the UMI position.

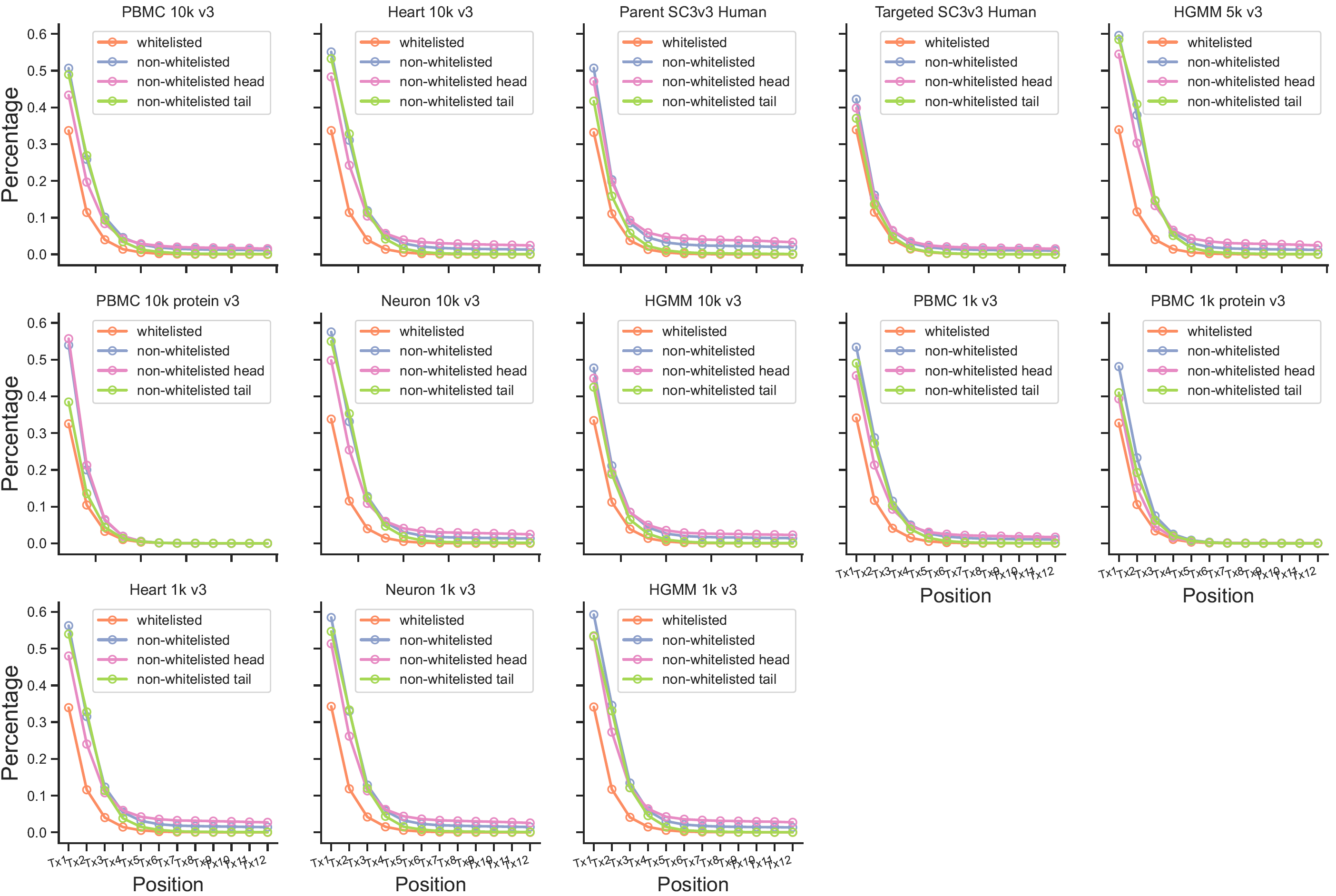

**Supplementary Figure 3**. Percentage of polyTs in UMIs on 13 datasets using chemistry V3 with respect to their barcodes that are or are not whitelisted or those non-whitelisted barcodes that have their head/tail regions overlapped with head/tail regions of barcodes in the whitelist. Each of the barcodes is split into two parts with the first 8 nucleotides denoted as the head region and the last 8 nucleotides denoted as the tail region. We observe the percentage of T increases towards the end of the UMI position.

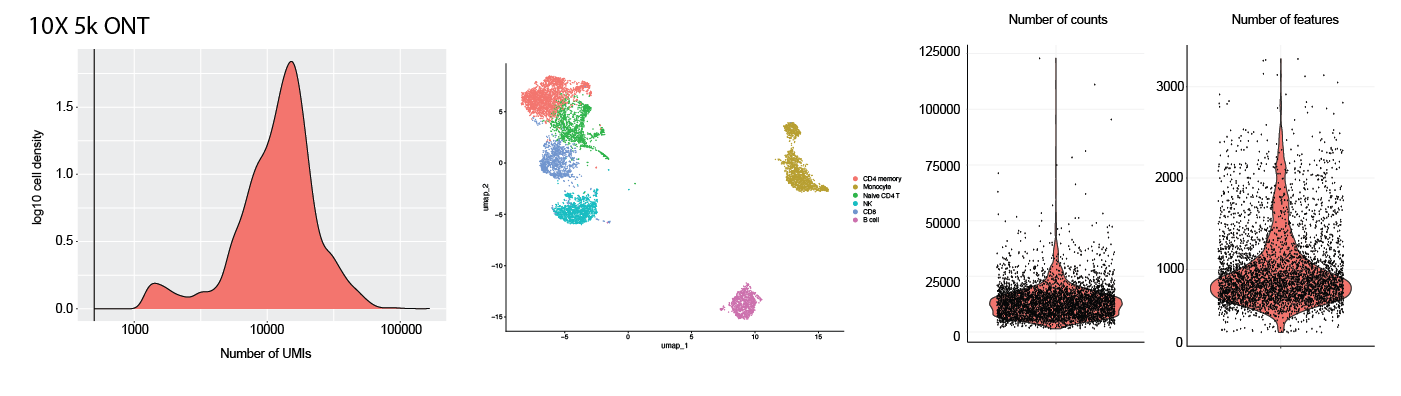

**Supplementary Figure 4:** 10X chromium quality control showing the 5k PBMC data. The data plotted on the left shows the number of UMIs captured. The middle plot shows the annotation of the cells and clusters. The right hand side shows the number of count and features for each cell plotted as a violin plot.

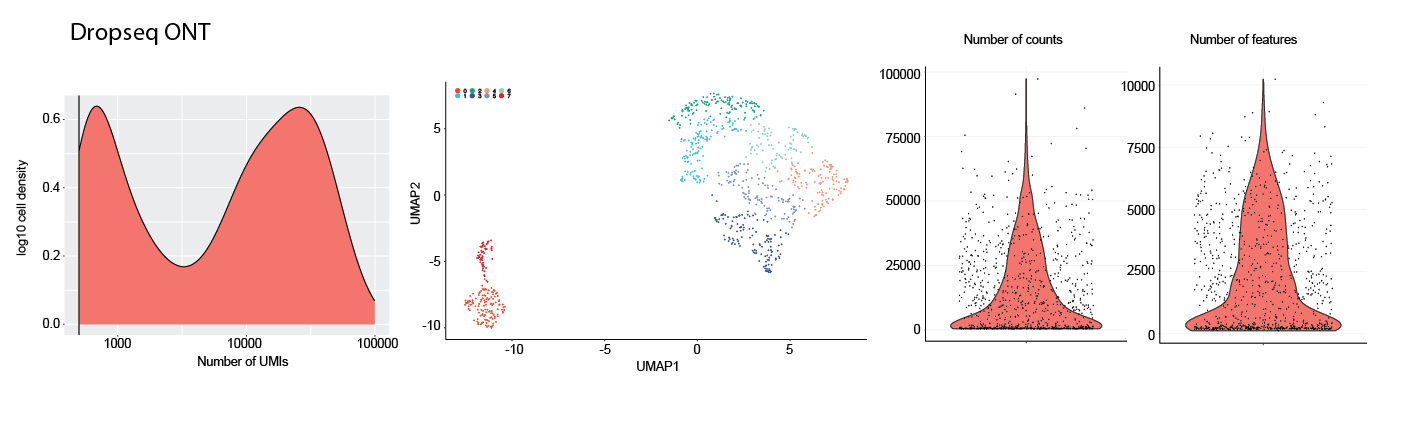

**Supplementary Figure 5:** Dropseq quality control showing the Makosko et al 2015 data. The data plotted on the left shows the number of UMIs captured. The middle plot shows the annotation of the cells and clusters. The right hand side shows the number of count and features for each cell plotted as a violin plot.

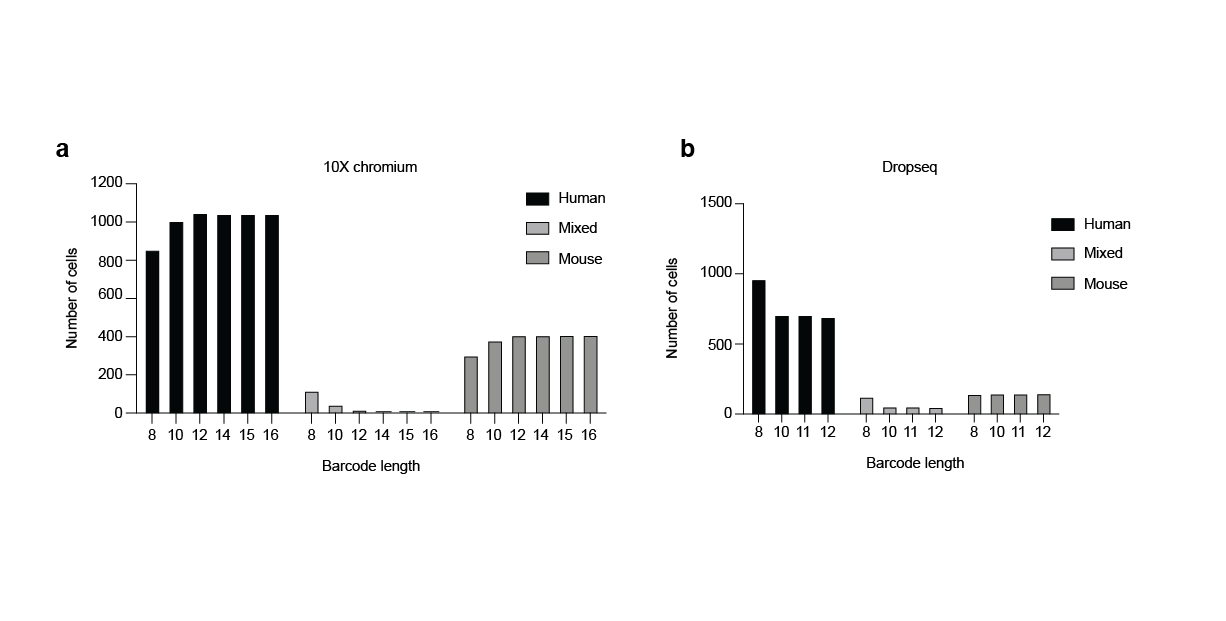

**Supplementary Figure 6**. Jurkat (70%) and 5TGM1 (30%) cells were co-encapsulated using both the 10X Chromium device and Dropseq, under the assumption that compromised barcode assignment would lead to a higher incidence of mixed cells. The expectation was that barcode truncation would result in a noticeable increase in mixed cell counts. However, analysis of both 10X Chromium and Dropseq datasets showed no significant rise in mixed cells, suggesting that synthesis irregularities have minimal impact on barcode assignment.

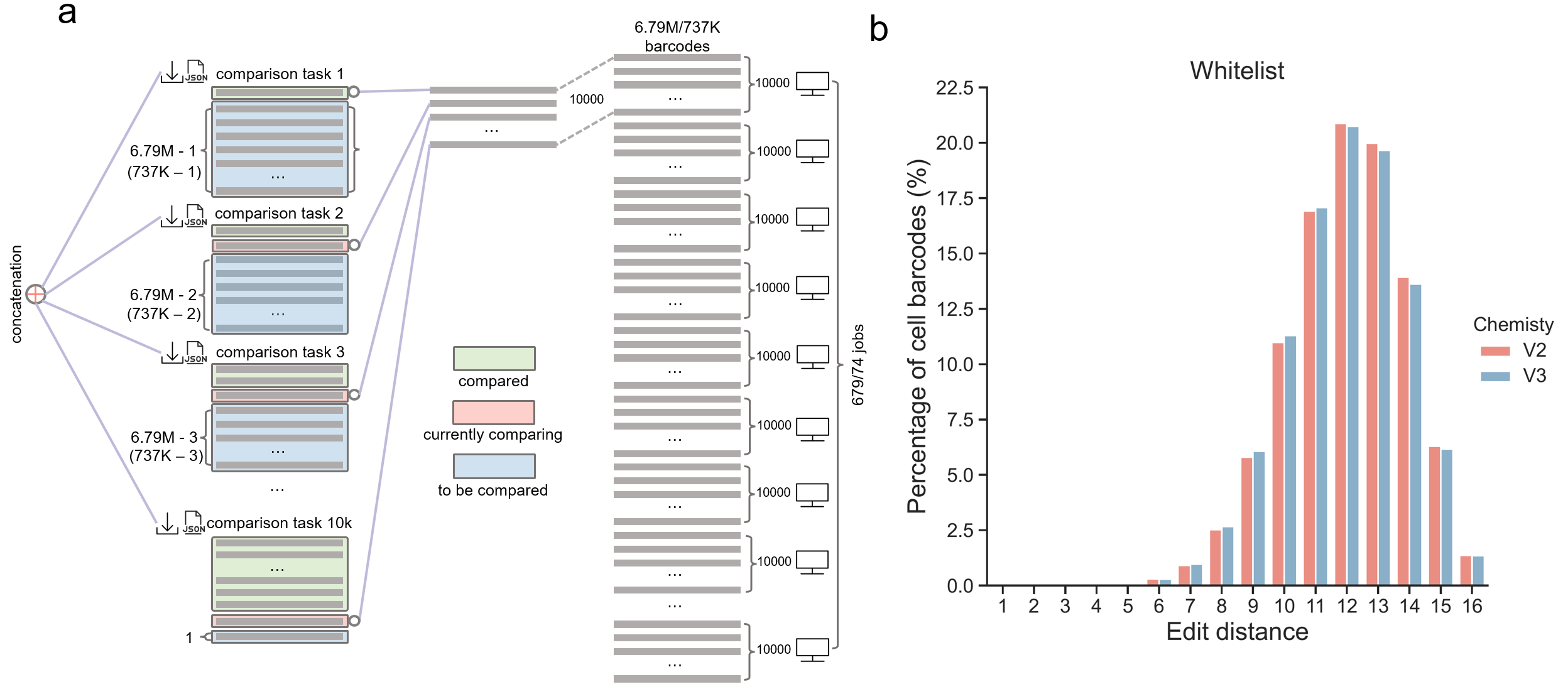

**Supplementary Figure 7**. All-against-all comparison of edit distances between cells barcodes from the whitelists of V2 (737K barcodes) or V3 chemistry (6.79M barcodes). a) In this study, we developed a distributed computing framework to expedite the computation of edit distances among a vast array of cell barcodes, specifically employing the whitelist associated with V3 chemistry as a reference case. The framework initiated 679 discrete computational jobs, each tasked with conducting comparisons between sets of 10,000 barcodes and those yet to be compared, resulting in 10,000 distinct comparison tasks within each job. For example, within the first job, the initial task computes the edit distances between the premier barcode and the remaining barcodes (totaling approximately 6.79 million minus one), and the 10,000th task calculates the edit distances between the 10,000th barcode and the subsequent barcodes (6.79 million minus 10,000). Sequentially, for the second job, the first task assesses the edit distances between the 10,001st barcode and the remaining set (6.79 million minus 10,001), indicating that barcodes within a designated green region are excluded from further comparisons. In this scheme, each task involves comparing a single barcode, delineated within a red region, against those within a blue region. The outcomes of these comparisons are documented in individual JSON files for each task. Upon completion of a job, the 10,000 resultant JSON files are amalgamated into a singular file. The computational process persists until the final task of the 679th job is concluded, thereby ensuring a systematic and non-redundant comparison across the entirety of the barcode dataset. This approach significantly conserves computational resources by reducing the total number of comparisons required to (6.79M×(6.79M-1))/2 in total. See main text for further explanation. b) Percentage of cell barcodes with respect to edit distances. Each bar represents the percentage of cell barcodes in a given edit distance away from the rest of the barcodes.

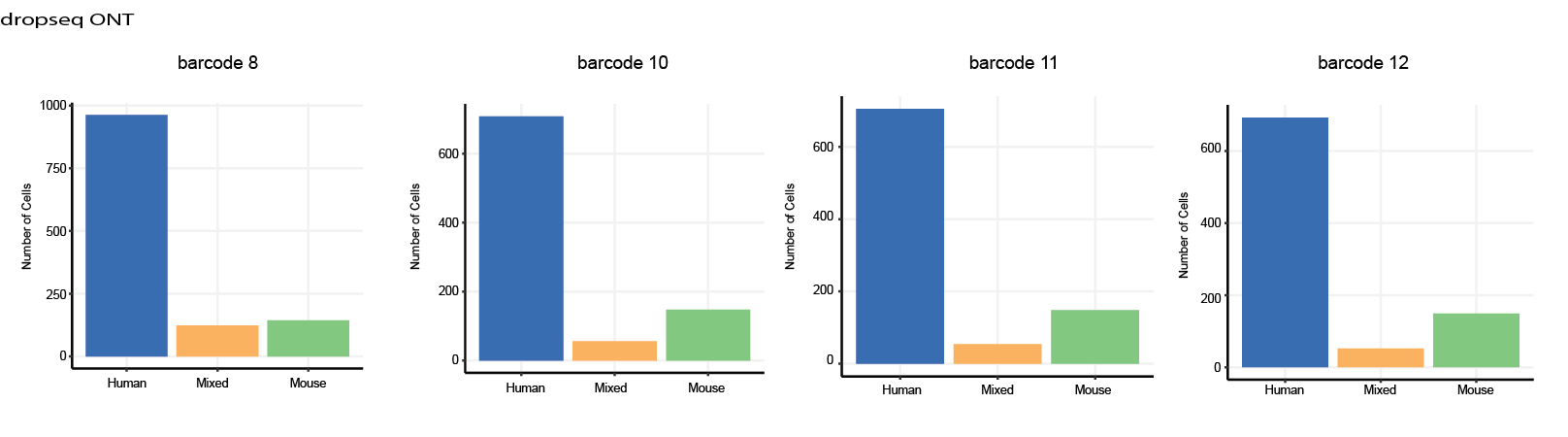

**Supplementary Figure 8**. Assessment of Barcode Length Reduction on Cell Calling Efficacy. This figure presents a systematic evaluation of the influence of incremental reductions in barcode sequence length (1, 2, 3, and 4 bases) on the proficiency of cell calling in a human and mouse mixing experiment. The analysis demonstrates that cell calling capacity remains largely unaffected by reductions in barcode length until a critical threshold of 8 bases is reached, beyond which the ability to accurately identify cells is compromised. This suggests that truncation issues seen with synthesis errors is likely not impacting the ability to assign reads to cells.

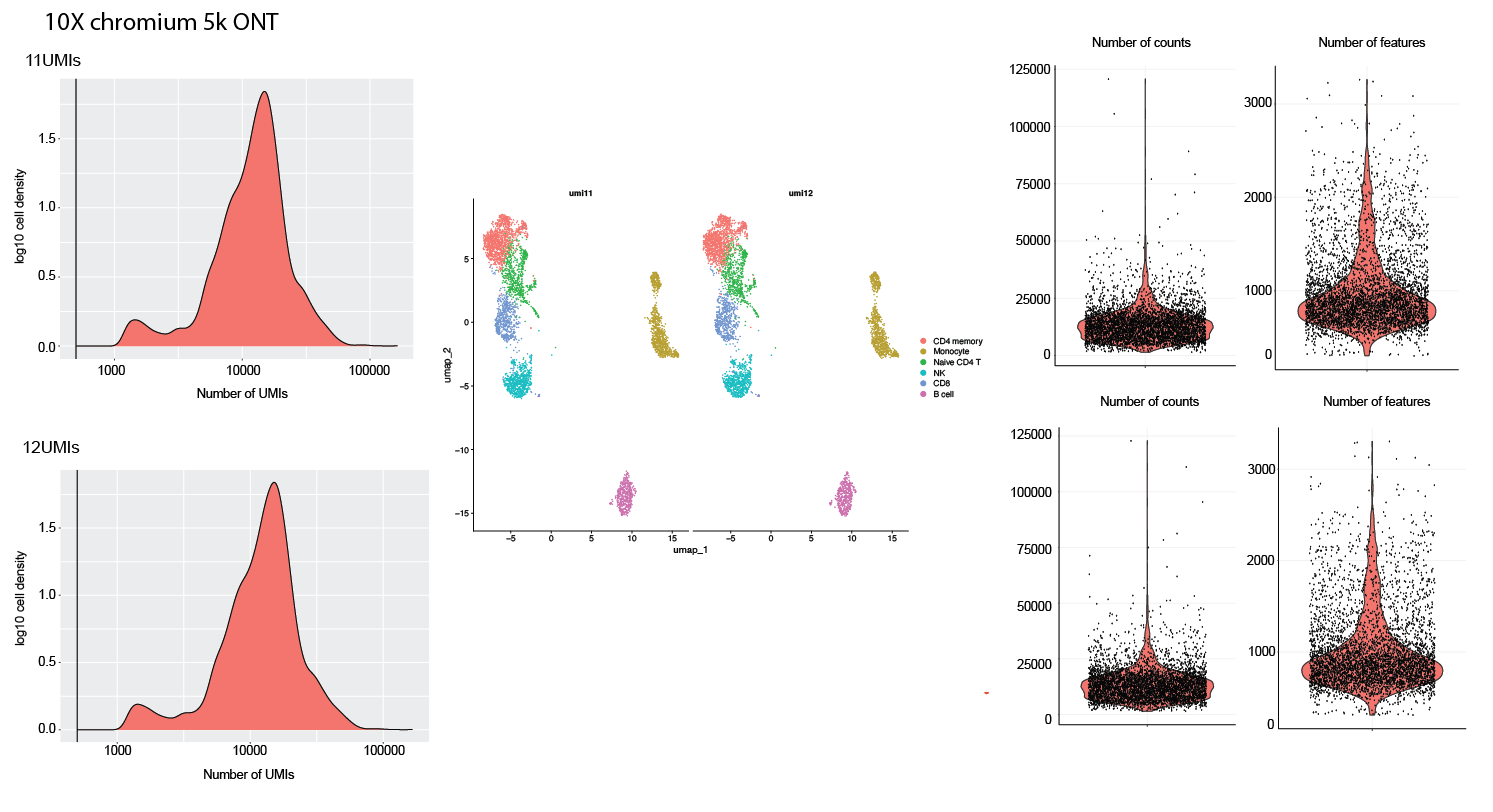

**Supplementary Figure 9:** 10X data showing quality plots showing the 5k 10X public data sequenced using ONT. The data plotted on the left shows the number of UMIs captured using 11and 12 UMI lengths. The middle plot shows the annotation of the cells and accompanies Fig. 2e and Fig. 2f. The right hand side shows the number of count and features for each cell plotted as a violin plot.

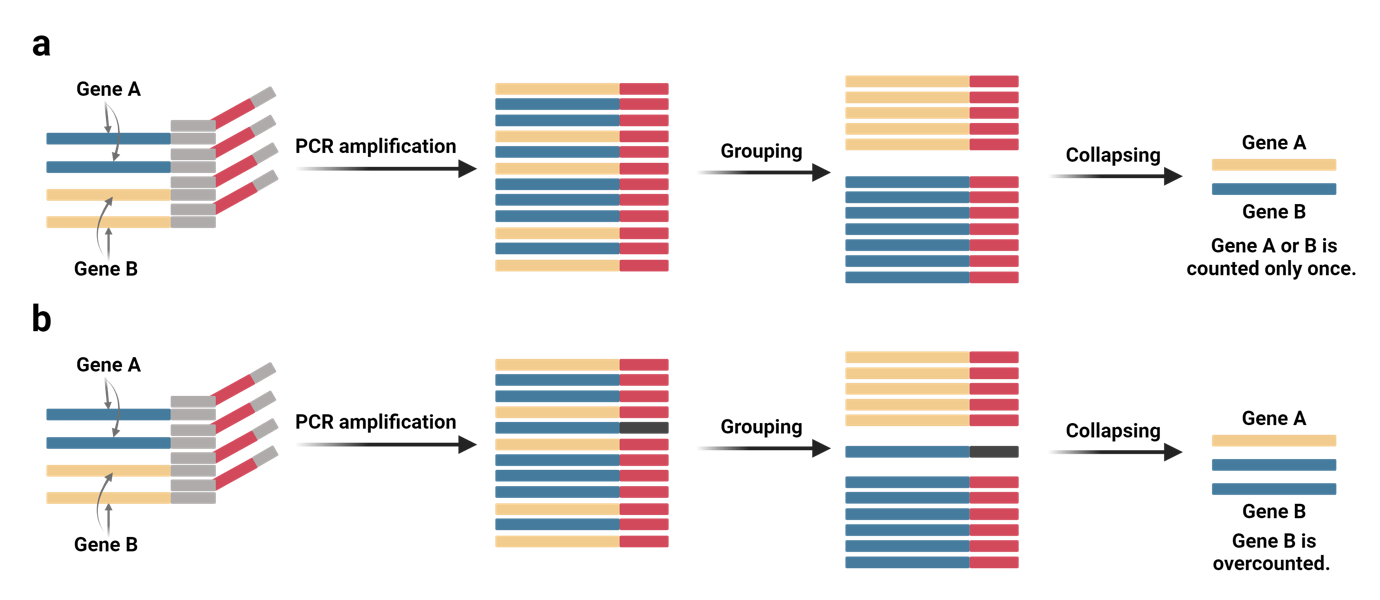

**Supplementary Figure 10:** Empirical evaluation of transcript counting with Common Molecular Identifiers (CMIs). **a**, An Ideal CMI collapsing example: In this scenario, transcripts (two blue and two green) are labelled with a common molecular identifier barcode (CMIs; labelled as red) and amplified via PCR. During transcripts grouping, all transcripts are labelled with the same common sequence. Therefore, following demultiplexing, each transcript should receive a count of one for every instance of detection. b, Increased counts result from the introduction of errors: This figure illustrates the effect of the errors within the CMI sequence. Any error introduced during PCR or sequencing creates a new CMI (labelled as yellow), resulting in an increase in transcript counts. This allows empirical evaluation of the effect of errors on the counting of transcripts, providing valuable insights into the accuracy of transcript quantification.

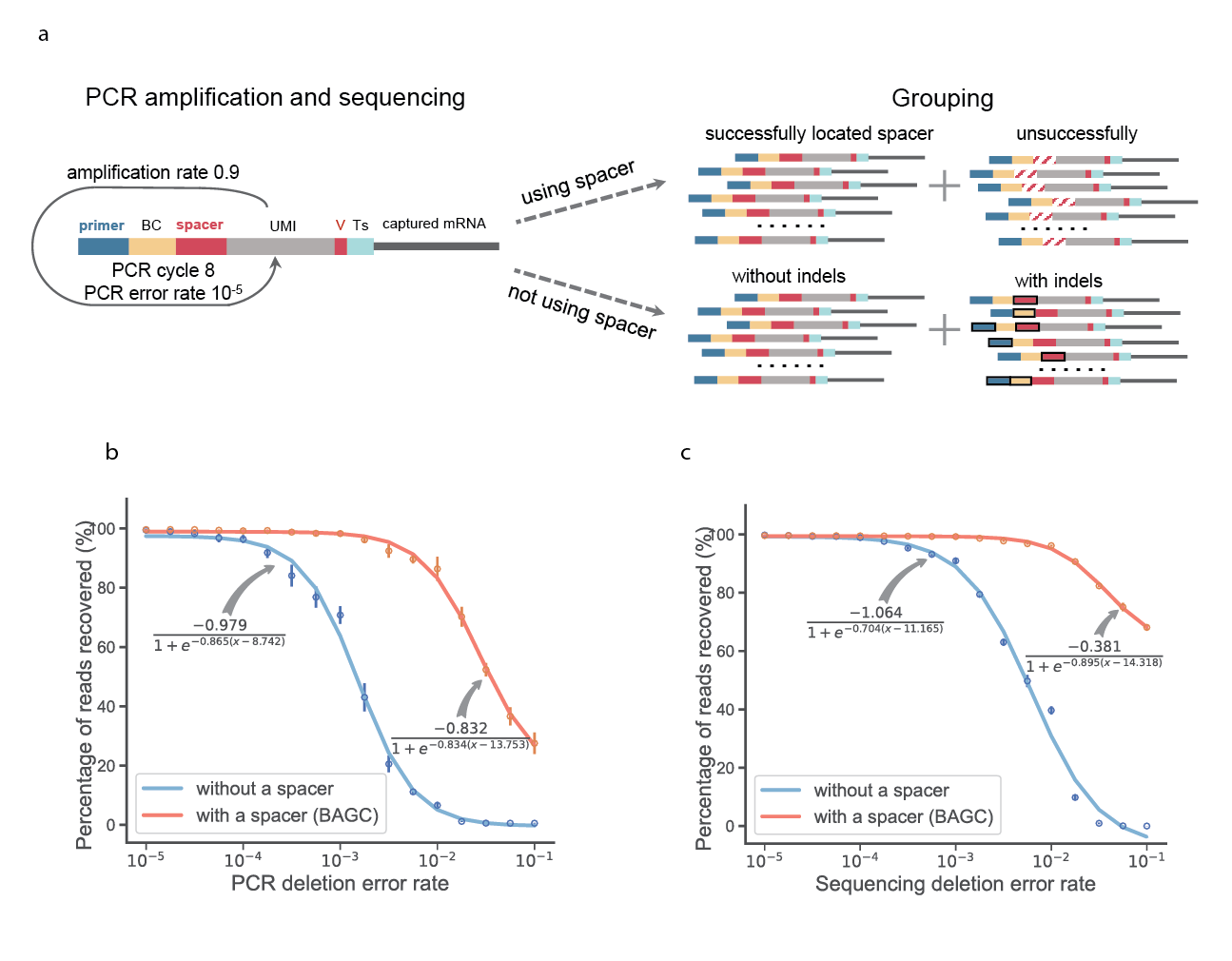

**Supplementary Figure 11:** Comparative Analysis of UMI Identification Efficacy with and without Anchor Utilisation. **a**) Illustrates the simulation workflow for assessing UMI (Unique Molecular Identifier) recognition efficiency employing an anchored versus non-anchored approach. Key features include the depiction of potential indels (insertions/deletions) or substitution errors within the anchor region (represented by white slashes) and other critical components such as primers, barcodes, or UMIs (denoted by dark squares). In simulation scenarios utilising an anchor, reads exhibiting errors in the anchor region are categorised as unsuccessful in UMI identification. Conversely, in the absence of an anchor, UMI identification success is determined by the direct analysis of the UMI sequence itself, with indels preceding the UMI indicating identification failure. **b** **and c**) show the outputs of simulating PCR and sequence error rates. The data illustrate the fidelity of UMI (Unique Molecular Identifier) identification across a spectrum of sequencing and PCR-induced deletion rates, providing a quantifiable measure of methodological resilience under diverse error profiles. This analysis not only benchmarks the accuracy of UMI detection but also underscores the robustness of the anchor-based identification process in the face of sequence deletions attributable to sequencing artifacts or PCR amplification errors. Methodological details and the analytical framework underpinning these evaluations are elaborated within the Methods section.

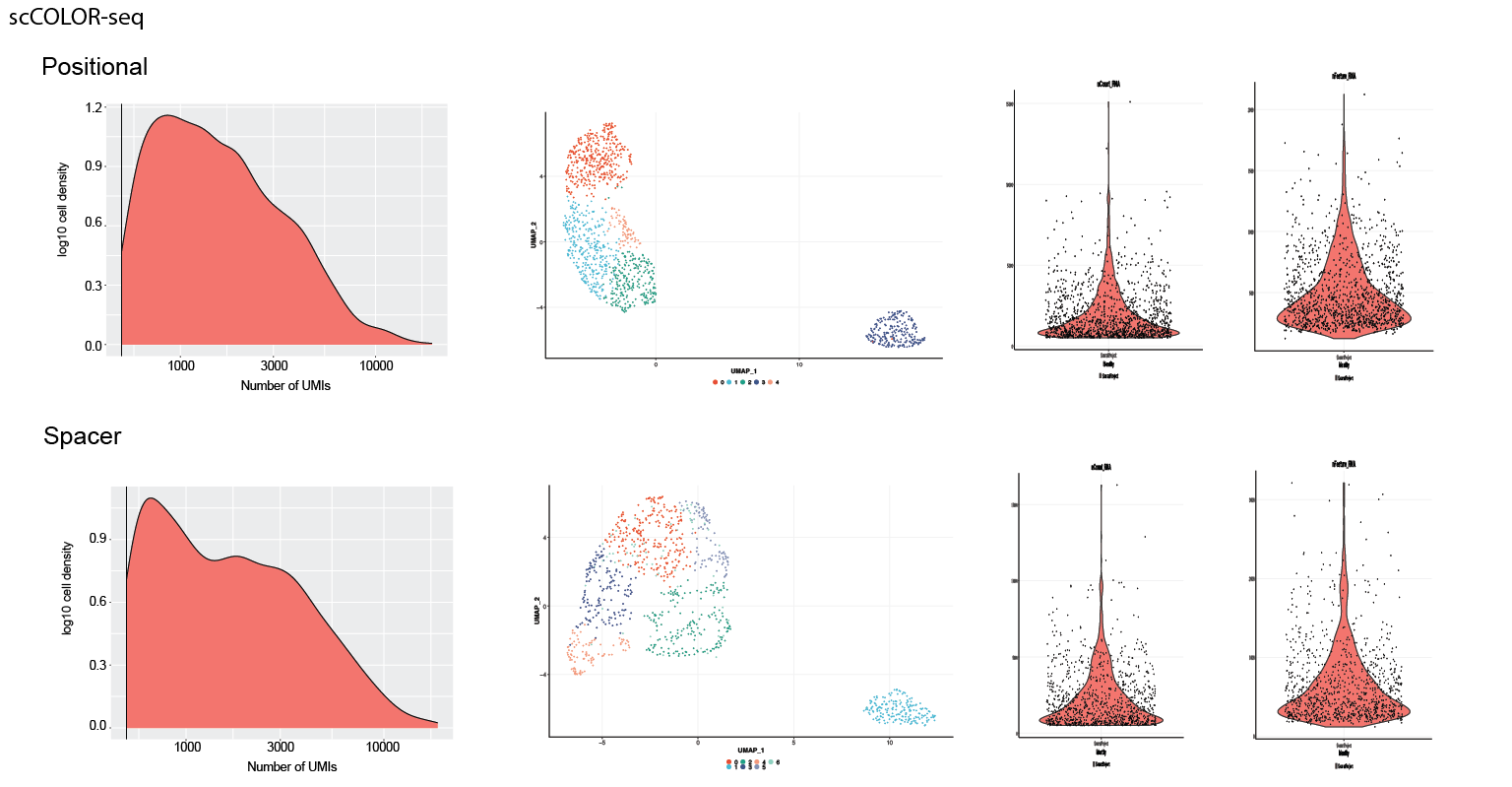

**Supplementary Figure 12:** scCOLOR-seq showing the quality metrics for the positional and anchor analysis approaches. The data plotted on the left shows the number of UMIs captured using positional and anchor approaches. The middle plot shows the annotation of the cells into clusters. The right-hand side shows the number of count and features for each cell plotted as a violin plot.
